# ANALYSIS OF FACTORS THAT REGULATE HIV-1 FUSION IN REVERSE

**DOI:** 10.1101/2025.03.10.642481

**Authors:** Ayna Alfadhli, Robin Lid Barklis, Fikadu G. Tafesse, Eric Barklis

**Affiliations:** Department of Molecular Microbiology and Immunology, Oregon Health and Sciences University, Portland, Oregon, 97239-3098, USA

**Keywords:** HIV-1, fusion, lipid, membranes

## Abstract

Based on observations that HIV-1 envelope (Env) proteins on the surfaces of cells have the capacity to fuse with neighboring cells or enveloped viruses that express CD4 receptors and CXCR4 co-receptors, we tested factors that affect the capacities of lentiviral vectors pseudotyped with CD4 and CXCR4 variants to infect Env-expressing cells. The process, which we refer to as fusion in reverse, involves the binding and activation of cellular Env proteins to fuse membranes with lentiviruses carrying CD4 and CXCR4 proteins. We have found that infection via fusion in reverse depends on cell surface Env levels, is inhibitable by an HIV-1-specific fusion inhibitor, and preferentially requires lentiviral pseudotyping with a glycosylphosphatidylinositol (GPI) anchored CD4 variant, and a cytoplasmic tail-truncated CXCR4 protein. We have demonstrated that latently HIV-1-infected cells can be specifically infected using this mechanism, and that activation of latently infected cells increases infection efficiency. The fusion in reverse approach allowed us to characterize how alteration of CD4 plus CXCR4 lipid membranes affected Env protein activities. In particular, we found that perturbation of membrane cholesterol levels did not affect Env activity. In contrast, viruses assembled in cells deficient for long chain sphingolipids showed increased infectivities, while viruses that incorporated a lipid scramblase were non-infectious. Our results yield new insights as to factors that influence envelope protein functions.

## INTRODUCTION

The trafficking of wild type (WT) HIV-1 envelope (Env) proteins to viral assembly sites at the plasma membranes (PMs) of infected cells is a complicated process that involves travel via a vesicular transport pathway, trimerization enroute, and processing in the Golgi by furin or a furin-like protease into their surface (SU; gp120) and transmembrane (TM; gp41) components (1–9). Evidence also indicates that because their cytoplasmic tails (CTs) carry YxxL and LL internalization signals, they can be internalized into endosomes where they may be shunted off for lysosomal degradation, or returned to the PM (1–9). The result of this is a reduced number of WT Env trimers at the PMs of infected cells, and surprisingly few WT Env proteins are incorporated into virus particles (1, 10–12). In this regard, it’s noteworthy that some Env variants with CT-truncations are present in PMs and virus particles at significantly higher levels than WT proteins, and one of these, which we refer to as 753Stop, still retains its YxxL internalization signal (11–13).

Once assembled into virions, Env proteins must operate in a lipid membrane environment that is considerably different than most PM regions (14–24). In addition to phosphatidylinositol-(4,5)-bisphosphate, the binding partner of the HIV-1 precursor Gag (PrGag) matrix (MA) domain, viral membranes are highly enriched in phosphatidylserine (PS), phosphatidylethanolamine (PE), saturated phosphatidylcholines (PCs), cholesterol, and sphingolipids (SLs), including ceramide (Cer), sphingomyelin (SM), and hexosylceramides (HEX) (14–16). The importance of this lipid composition is underscored by the effects of tampering with it. In particular, we’ve shown that production of viruses in ceramide synthase 2 knockout (CerS2-/-) cells, which reduces the levels of long chain SLs, impairs the fusion functions of both WT and CT deletion (ΔCT) Env proteins (25, 84). We and others also have demonstrated that treatment of HIV-1 virions with the cholesterol binding compound, amphotericin B methyl ester (AME), dramatically reduces the fusion activity of WT, but not ΔCT viruses (13, 26–28). Finally, the Env fusion function also is dependent on the membrane leaflet lipid asymmetry in virions, and scramblase incorporation into HIV-1 virions blocks viral infection (29–34).

Interestingly, the Env receptor-binding and fusion functions can be active in the PMs of Env-expressing cells (35–39). Notably, when Env-expressing cells are in the presence of cells that carry the HIV-1 Env CD4 receptor plus a co-receptor (CXCR4 or CCR5), the combination can lead to cell-cell fusion and syncytia formation (35–39). Similarly, enveloped viruses carrying CD4 and a co-receptor have been shown to infect Env-expressing cells in a process we refer to as fusion in reverse (40–47). Recently, using cells expressing a different viral surface protein, the Spike protein of SARS-CoV-2, we demonstrated that lentiviruses decorated with a Spike-binding nanobody could specifically infect SARS-CoV-2-infected cells via a fusion in reverse mechanism (48). Based on these results, we reasoned that might be possible to infect HIV-1 Env-expressing cells in a related fashion.

We now report the ability to specifically infect HIV-1 Env-expressing cells with lentiviral vectors via a fusion in reverse mechanism. The optimized vectors utilize HIV-1 Gag and Pol proteins, and employ a C-terminally truncated CXCR4 protein, as well as a CD4 variant that is anchored to membranes by virtue of glycosylphosphatidylinositol (GPI) tail (48–51). The lentiviruses preferentially infect HIV-1 infected cells, or cells independently expressing HIV-1 Env, and infection efficiency is dependent on Env surface expression levels. Using this fusion in reverse approach, it was possible to assess how lipid variations in membranes containing the HIV-1 receptor and co-receptor affected the HIV-1 Env activity. Interestingly, we found that reducing the leaflet asymmetry of viral membranes containing receptors plus co-receptors effectively inhibited infection, that reduction of long chain SLs increased infectivity, and that AME treatment of receptor-containing membranes did not impair the activity of PM Env. We believe this system will be useful in the analysis of other factors involved in Env-mediated membrane fusion.

## RESULTS

### Analysis of Env-mediated cell-cell fusion

We recently observed that HIV-1-derived lentivirus vectors decorated with a nanobody against the SARS-CoV-2 Spike protein selectively infected SARS-CoV-2-infected cells via a fusion in reverse mechanism in which nanobody-Spike binding activated the fusion function of Spike (48). These results were reminiscent of efforts to infect HIV-1 Env-expressing cells with membrane-enveloped viruses carrying receptors and co-receptors (40–47). Based on this, we reasoned that it might be possible to employ a fusion in reverse approach to examine factors in target membranes that affect HIV-1 Env functions. As a preliminary test, we examined the capacities of Env, receptor and co-receptor combinations to function in cell-cell fusion assays. The method employed (Figure 1A) is similar to one previously described (52), and takes advantage of split green fluorescent protein (GFP) fragments to assemble into active fluorescent proteins (52–53). As shown (Figure 1A), one set of cells was cotransfected with an Env expression plasmid plus a PM-targeted large helix 1-10 GFP fragment, while a second set of cells was cotransfected with CD4 receptor-CXCR4 co-receptor pairs plus a PM-targeted small (helix 11) GFP fragment. After mixing the cells, fusion was assayed via the appearance of GFP-positive (GFP+) syncytia.

**Figure 1.**
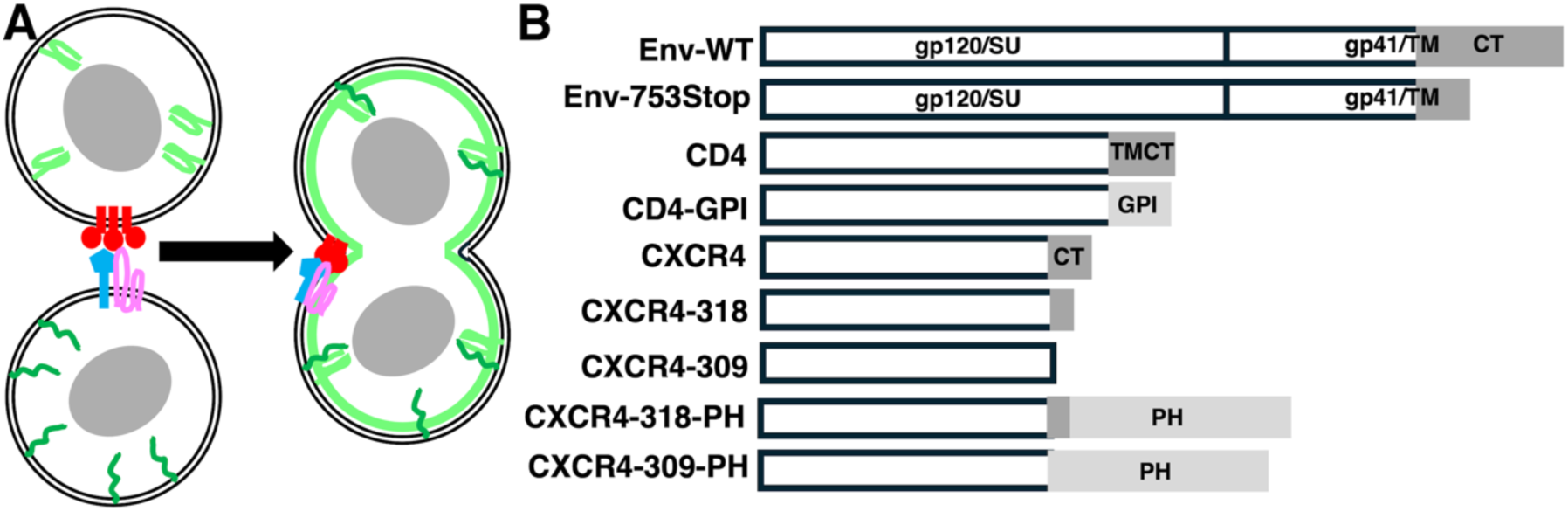
Cell-cell fusion assays and recombinant envelope and receptor constructs. **A.** Cell-cell fusion assays involve transfection of one set of cells with an Env expression construct (in red) plus a PM-targeted split GFP helix 1-10 expression construct (pcDNA3.1-PLCdeltaPH-GFP-H1-10; green), and transfection of a separate set of cells with expression vectors for CD4 (blue) and CXCR4 (pink) variants, plus a PM-targeted split GFP helix 11 expression construct (For pGFP-H11-PLCdeltaPH; green). One day post-transfection cells are seeded together onto coverslips, and scored for GFP+ syncytia two days later. B. HIV-1 Env expression constructs included one that produces the WT HIV-1 NL4-3 envelope protein (Env-WT: pMDG-NL43-Env-WT or SVIII-Env-WT) and one that produces an Env protein (Env-753stop: pMDG-NL43-Env-753stop), truncated at Env residue 753, before the CT amphipathic helices. CD4 expression constructs included one that expresses a Myc-tagged WT human CD4 protein (CD4: pCAGGS-CD4-Myc) or a variant in which the CD4 transmembrane and cytoplasmic tail (TMCT) domains were replaced with a GPI anchor (CD4-GPI: pCAGGS-CD4-GPI). CXCR4 expression constructs included a WT human CXCR4 expression construct (CXCR4: pcDNA3-CXCR4), constructs with partial (CXCR4-318: pcDNA3-CXCR4-318Stop) or complete (CXCR4-309: pcDNA3-CXCR4-309Stop) CXCR4 cytoplasmic tail (CT) deletions, and constructs in which CT truncations were fused to the PLCδPH that preferentially binds to PI(4,5)P2 (CXCR4-318-PH and CXCR4-309-PH).

For our initial purposes, an expression vector for the WT Env (Figure 1B, Env-WT) protein was used, along with a vector for the 753Stop Env protein (Env-753Stop), which carries a CT truncation, behaves similarly to a complete ΔCT Env protein, and which is incorporated at high levels into HIV-1 particles (11–13). Our CD4 vectors expressed a WT variant (Figure 1B, CD4) or a GPI-anchored CD4 (CD4-GPI), since GPI-anchored CD4 previously was shown to function as an HIV-1 receptor, and because GPI-anchored proteins may be assembled efficiently into HIV-1 particles (40–51). For co-receptors, we tested WT CXCR4 (CXCR4), which has been shown to be delivered, under some circumstances, to exosomes and enveloped viruses; and a C-terminally truncated variant (CXCR4-318), to remove residues involved in phosphorylation and internalization (40–47, 54–58).

Examples of our cell-cell fusion assays are shown in Figure 2A-F. In each case, one set of fusion-partner cells expressed the helix 11 GFP fragment plus CD4-GPI and CXCR4-318, while the other sets of cells expressed the large GFP fragment plus Env-WT (A-B), Env-753Stop (C-D), or no Env (E-F). Cells were monitored in parallel for nuclei using 4’,6-diamdino-2-phenylindole (DAPI) stain (A, C, E) or for GFP+ syncytia (B, D, F). Not surprisingly, few if any GFP+ syncytia were observed for the no Env incubation (Figure 2F), whereas some were seen with Env-WT (Figure 2B), and syncytia were numerous in the Env-753Stop incubation (Figure 2D). These observations were borne out when syncytia were quantified (Figure 2G). Notably, numbers of Env-WT syncytia were 20-fold higher than background, and Env-753Stop syncytia numbers were 10-fold higher than those for Env-WT. It also was apparent that all of the receptor plus co-receptor combinations yielded similar results.

**Figure 2.**
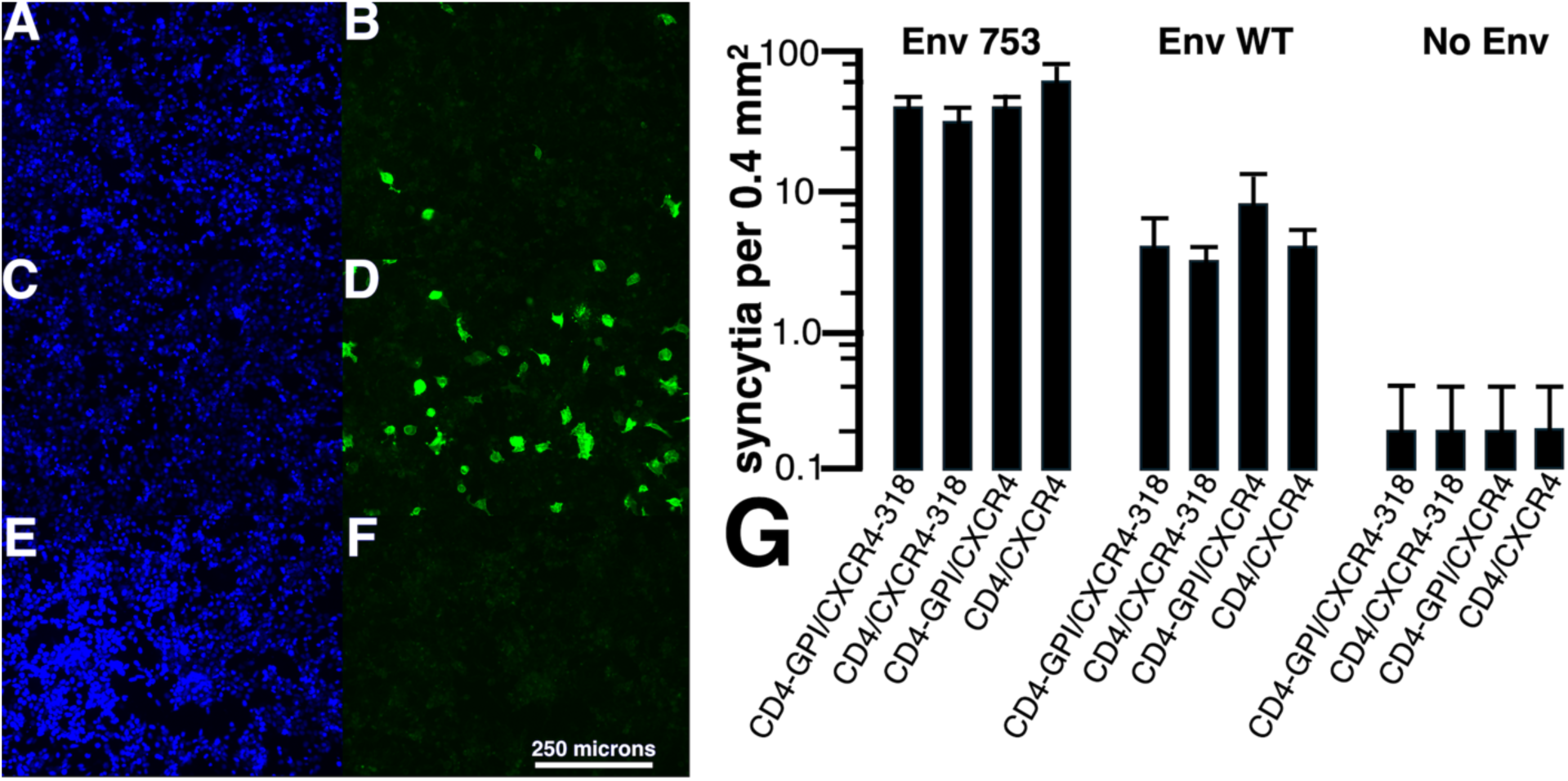
HIV-1 Env cell-cell fusion assays. Cell-cell fusion assays were performed as illustrated in Figure 1. Panels **A-F** show coincubations of cells transfected with pGFP-H11-PLCdeltaPH plus CD4-GPI plus CXCR4-318 mixed with cells transfected with pcDNA3.1-PLCdeltaPH-GFP-H1-10 plus either Env-WT (**A-B**), Env-753Stop (**C-D**) or a mock (**E-F**). Panels **A** and **B**, **C** and **D**, and **E** and **F** are matched photos imaged for DAPI nuclear stain (**A, C, E**) and GFP (**B, D, F**). Note that the size bar (panel **F**) pertains to all fluorescent images (**A-F**). In panel **G**, numbers of syncytia per 0.4 mm^2^ are averaged with standard deviations from five images each of coincubations of cells cotransfected split GFP constructs plus the indicated Env or CD4 plus CXCR4 expression construct variants. Note that numbers of syncytia for Env-753 were approximately ten-fold higher than syncytia for Env-WT, and that syncytia for Env-WT were approximately twenty-fold higher than syncytia for the Env-minus (No Env) control, and that differences between the sets were highly significant (P<0.001).

What might account for the 10-fold difference in syncytia formation with the Env-753Stop versus Env-WT proteins? Because an HIV-1 Env protein with a truncation similar to Env-753Stop had been selected for enhanced cell surface expression (11–12), we speculated that our observed increase in syncytia formation with Env-753Stop might stem from a high level of surface expression. To test this possibility, target cells expressing Env-WT versus Env-753Stop proteins were subjected to immunofluorescent detection of cell surface Env. Although we didn’t observe discernable differences in localization patterns of the different Env proteins (Figures 3A-B), it was noteworthy that Env imaging required higher exposure times for Env-WT versus Env-753Stop (67 ms in Figure 3B versus 10 ms in Figure 3A). To quantify our observations, we measured percentages of fluorescent cells (Figure 3C) and relative brightness levels (Figure 3D) of the Envs with two different anti-gp120(SU) antibodies (VRC3, 2G12). Both antibodies detected moderately lower (2.5-fold) percentages of Env-WT fluorescent cells, but substantial reductions (10-fold) in Env-WT brightness levels that correlate with surface expression levels. Given that differences in Env-WT and Env-753Stop surface expression levels matched the differences we observed in cell-cell fusion capacities (Figure 2), we conclude that the increased frequency of Env-753Stop-mediated fusion derives from its high surface availability.

**Figure 3.**
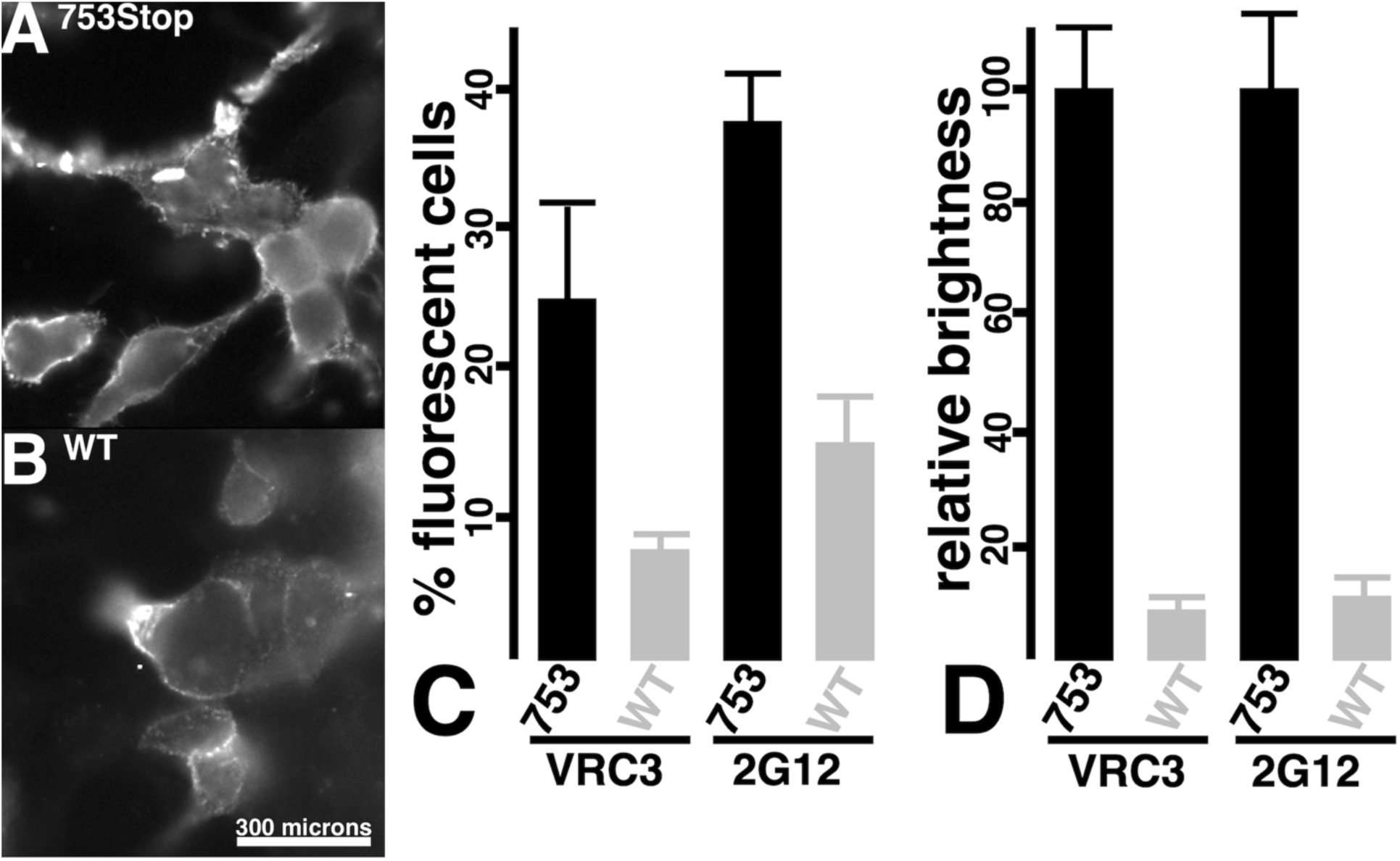
Cell surface expression of Env proteins. **A-B.** Cells transfected with the indicated Env expression plasmids were subjected to indirect immunofluorescent detection of surface Env proteins in the absence of cell permeabilization with the human 2G12 anti-Env antibody as a primary antibody, and a fluorescent anti-human IgG secondary antibody. Note that the size bar in panel **B** pertains to both panels, and that the exposure times for panels **A** and **B** respectively were 10 and 67 ms. Panels **C** and **D** respectively indicate percentages of fluorescent cells and relative brightness values for fluorescent cells, using the cells transfected with the indicated Env expression constructs and either VRC3 or 2G12 anti-Env antibodies for detection. Averages and standard deviations were obtained from a minimum of nine images for panel **C**, a minimum of 14 cells for panel **D** (VRC3), and a minimum of 36 cells for panel **D** (2G12). Differences between all pairs were highly significant (P<0.001).

### Viral infection via fusion in reverse

Given that CD4-GPI and CXCR4-318 seemed likely candidates to be incorporated into HIV-1 particles (48–51, 54–58), and that they efficiently mediated cell-cell fusion reactions (Figure 2), we tested their propensity for targeting the infection of Env-expressing cells. To do so, viruses produced by cotransfection of cells with psPAX2, pLVX-puro-Xho-ATG-βGal, CD4-GPI and CXCR4-318 plasmids were used to infect Env-expressing cells (Figure 4A), scoring for infection by βGal expression. As shown in Figure 4B, our results clearly showed that the CD4-GPI plus CXCR4-318 pseudotyped viruses specifically infected Env-expressing cells. In particular, infection of Env-WT cells was over ten times greater than Env-minus (mock) cells, while infection of Env-753Stop cells appeared ten times more efficient than infection of the Env-WT cells. These results closely mirrored our cell-cell fusion results (Figure 2), implying that infection efficiency also is dependent on cell surface Env protein levels (Figure 3). To verify that infections were dependent on the Env fusion function, separate infections were performed in the absence or presence of the HIV-1 Env fusion inhibitor T-20 (Enfuvirtide; 59-61). As depicted in Figure 4C, T-20 almost completely abolished infectivity, supporting the interpretation that infection occurred by a fusion in reverse process.

**Figure 4.**
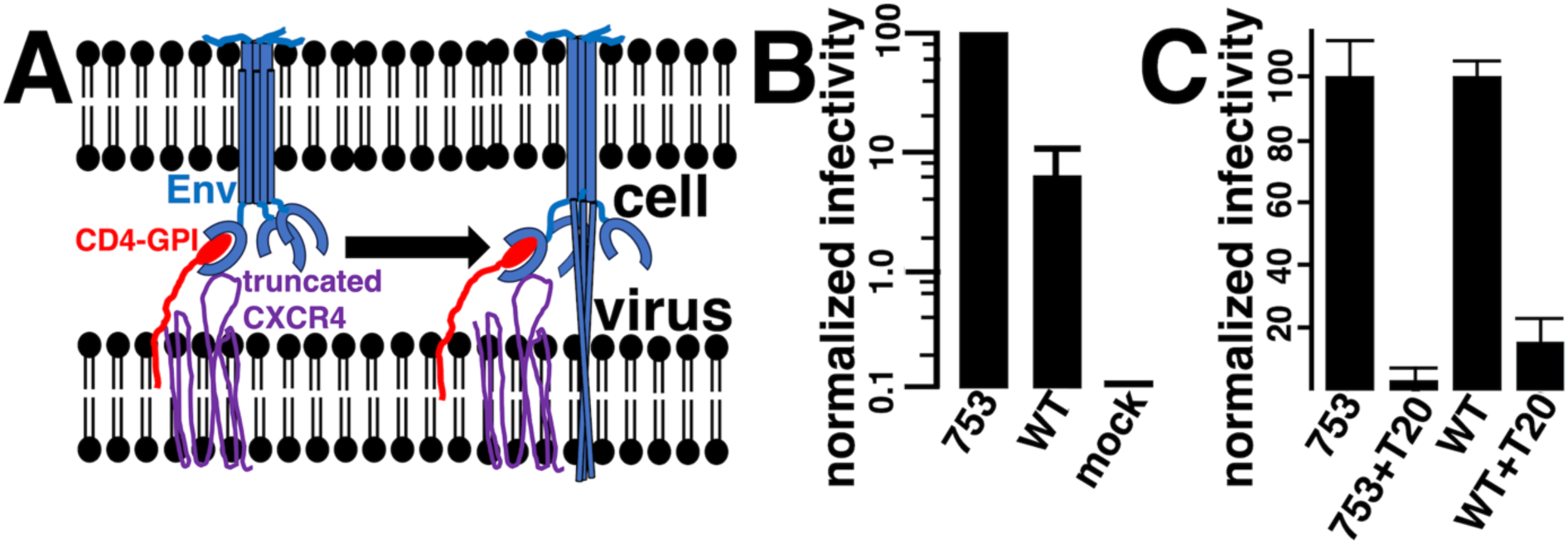
Infection of Env-expressing cells via fusion in reverse. **(A)** Target cells expressing different Env variants are infected with a βGal-transducing, HIV-1-based, lentivirus vector pseudotyped with a GPI-anchored CD4 variant (CD4-GPI) plus a C-terminally-truncated CXCR4 protein (CXCR4-318). Receptor plus co-receptor binding to cellularly expressed Env activate the Env fusion machinery resulting in virus infection. **(B)** Shown are βGal activity-derived infectivities of cells expressing Env-753Stop, Env-WT, or no Env (mock), normalized to the Env-753Stop cells. Note that results are averaged from five independent experiments each, are plotted on a log scale graph to facilitate comparison, and that standard deviations for the Env-753Stop and mock cells were too small to be observed on the graph. **(C)** Cells expressing the indicated Env proteins were infected with CD4-GPI plus CXCR4-318 pseudotyped bGal-transducing HIV-1 lentivirus vectors in the absence (753, WT) or presence (753+T20, WT+T20) of HIV-1 Env fusion inhibitor T20. Results for pairs of infections are normalized to the infectivities of viruses in the absence of T20. Averages and standard deviations derive from three separate infections, and differences between 753 and WT, and WT and mock were highly significant (P<0.001), as were differences between untreated and treated samples.

To help optimize the receptor and co-receptor requirements for infection, different CD4 and CXCR4 combinations (Figure 1B) were tested for their capacities to infect cells expressing Env-753Stop (Figure 5A) and Env-WT (Figure 5B), and normalized to results of the CD4-GPI (**gpi**) plus CXCR4-318 (**318**). As expected, little or no infection was observed when either receptor or co-receptor was used alone (**gpi/-; -/318)**. Surprisingly, substitution of WT CD4 for CD4-GPI also dramatically reduced infection efficiency (**CD4/318**). In contrast, replacement of CXCR4-318 with WT CXCR4, or with CXCR4 carry a nearly complete CT truncation (CXCR4-309; **309**), or with CXCR4 variants bearing C-terminal fusions to a PI(4,5)P2-binding pleckstrin homology (**PH**; 62-64) domain (CXCR4-318PH [**318PH**], CXCR4-309PH [**309PH**]) had little effect on infectivities. Nevertheless, the most efficient combination for infection either Env-753Stop or Env-WT cells remained the CD4-GPI plus CXCR4-318 combination.

**Figure 5.**
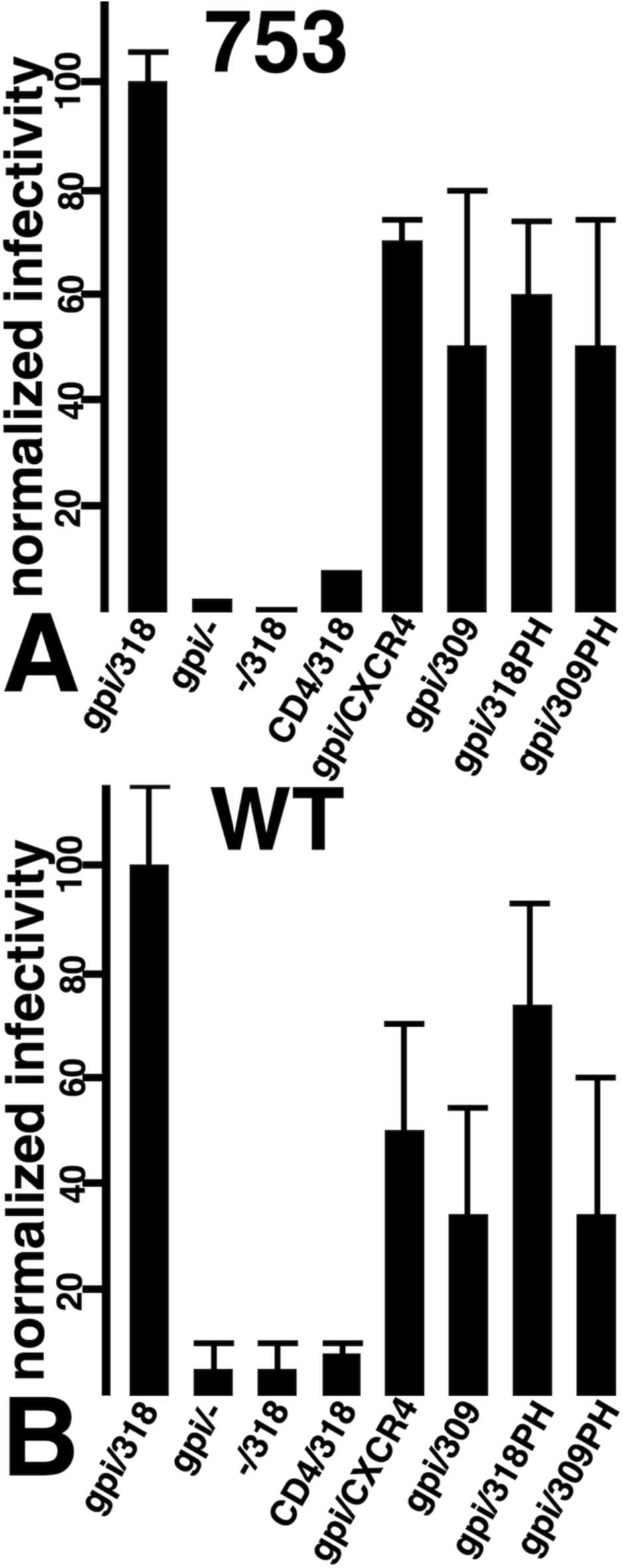
Fusion in reverse infectivities for different Env, receptor, and co-receptor combinations. Cells expressing the 753Stop **(A)** or WT HIV-1 Env **(B)** were infected with βgal transducing HIV-1 particles carrying the indicated receptor plus co-receptor combinations (CD4, CD4-GPI [gpi], - [mock], CXCR4, 318 [CXCR4-318], 309 [CXCR4-309], 318PH [CXCR4-318-PH], 309PH [CXCR4-309-PH]). At 72 h post-infection, cells were subjected to βgal assays and βgal activities are plotted as infectivities relative to viruses carrying the CD4-GPI/CXCR4-318 receptor/co-receptor combinations. Averages and standard deviations derive from the following numbers of experiments: 753/gpi/318, 9; 753/gpi/-, 2; 753/-/318, 2; 753/CD4/325, 2; 753/gpi/CXCR4, 4; 753/gpi/309, 3; 753/gpi/318PH, 5; 753/gpi/309PH, 5; WT/gpi/318, 9; WT/gpi/-, 2; WT/-/318, 2; WT/CD4/318, 3; WT/gpi/CXCR4, 6; WT/gpi/309, 4; WT/gpi/318PH, 5; WT/gpi/309PH, 5. Note that differences between the gpi/318 samples versus the gpi/-, -/318, and CD4/318 samples had probabilities of P<0.001.

Because none of the CXCR4 variants gave dramatically different infectivity results (Figure 5), and because our anti-CXCR4 antibodies were ineffective in immunoblotting, we did not examine CXCR4 variant virion-incorporation. Nevertheless, all the CXCR4 gave similar cellular immunofluorescence staining patterns, suggesting similar expression levels. However, because of the significant disparity between cell-cell fusion (Figure 2) and fusion in reverse infection results (Figure 5) for WT CD4, it was worthwhile to examine CD4 and CD4-GPI virion incorporation levels in greater detail. Initially, virions from cells expressing HIV-1 Gag and GagPol plus either CD4 or CD4-GPI were collected by sedimentation through 20% sucrose cushions, and subjected to immunoblot detection of PrGag, CA, and CD4 proteins. As shown in Figure 6A, CD4 migrated, as expected, at a slightly slower mobility than CD4-GPI, and the observed viral CD4-to-CA ratio appeared greater than that of CD4-GPI. As a more stringent test, the two CD4 variants were expressed in the presence or absence of Gag plus GagPol proteins, and media samples were sedimented through 30% sucrose cushions to reduce possible exosome or membrane vesicle contamination. Cell and viral samples so prepared were processed for immunodetection with anti-CA (Figure 6B) and anti-CD4 (Figure 6C) antibodies. Our anti-CA immunoblots gave the expected PrGag and CA bands in samples derived from cells transfected with the psPAX2 (psPAX) Gag and GagPol expression construct. The CD4 and CD4-GPI proteins also were readily detected in cellular samples (Figure 6C, top panel). Importantly, both WT CD4 and CD4-GPI were present in virus samples from cells cotransfected with psPAX2, while only trace amounts were present in samples processed similarly from cells not expressing Gag and GagPol proteins. Based on virus-to-cell CD4 ratios, it appeared that CD4-GPI might be preferentially incorporated into virions, consistent with observations with other GPI-tagged proteins (48). However, when we calculated viral CD4-to-CA ratios the WT CD4 and CD4-GPI protein values were within 10% of each other. These results agree with studies that have demonstrated CD4 incorporation into HIV-1 virions (42, 44–45). Nevertheless, our observations strongly suggest that in the context of HIV-1 viral membranes, CD4-GPI is considerably more competent than CD4 in fostering Env-mediated fusion.

**Figure 6.**
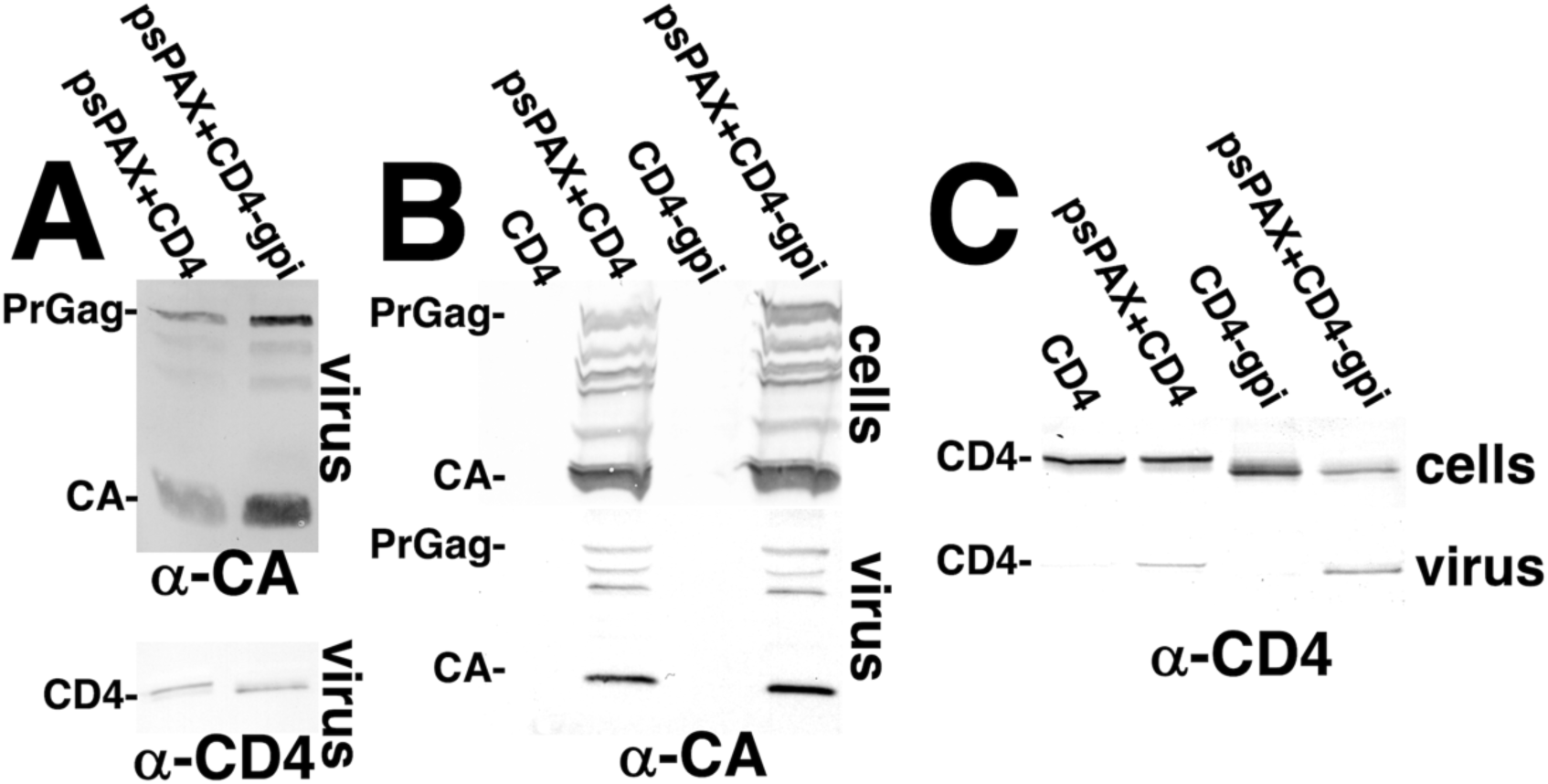
CD4 incorporation into virions. **A.** Media samples from cells transfected with psPAX2 (psPAX) plus the CD4 or CD4-gpi expression constructs were pelleted through 20% sucrose cushions, separated by electrophoresis and immunoblotted for detection of CA and CD4 immunoreactive proteins. PrGag, CA and CD4 proteins were identified based on their immunoreactivities and migration mobilities relative to protein molecular weight standards run in parallel. **B-C.** Cell samples (top panels) and media samples that were pelleted through 30% sucrose cushions (bottom panels) were collected from cells transfected with the indicated constructs, separated by electrophoresis, and immunoblotted for detection of PrGag and CA **(B)** or CD4 and CD4-gpi **(C)**. Note that viral CD4-to-CA and CD4-gpi-to-CA ratios were within 10% of each other.

### Targeted transduction of HIV-1 infected cells

Beyond demonstrating that cells transfected to express HIV-1 Env could be targeted for infection, it was of interest to verify that HIV-1-infected cells could be similarly targeted. For these experiments, we employed the HIV-1-infected J1.1 Jurkat T cell line that has proven useful in latency reactivation studies, and can be induced by treatment with tumor necrosis factor alpha (TNFα; 65-69). As expected, and as illustrated in Figure 7, TNFα treatment of J1.1 cells increased cellular levels of both Gag (Figure 7A; PrGag, CA) and Env (Figure 7B; gp160, gp41) proteins. Thus, for our purposes, we tested the propensity of CD4-GPI plus CXCR4-318 pseudotyped lentivirus vectors to transduce Jurkat, J1.1, and TNFα-induced J1.1 (J1.1+TNFα) cells. As illustrated in Figure 7C, relative to background transduction of Jurkat cells, βGal delivery to J1.1 cells via fusion in reverse was 7-fold higher, and infection levels rose to 10-fold over background with induced J1.1 recipients. These results are consistent with previous results in showing that the fusion in reverse mechanism can be utilized to target HIV-1-infected cells (40–47).

**Figure 7.**
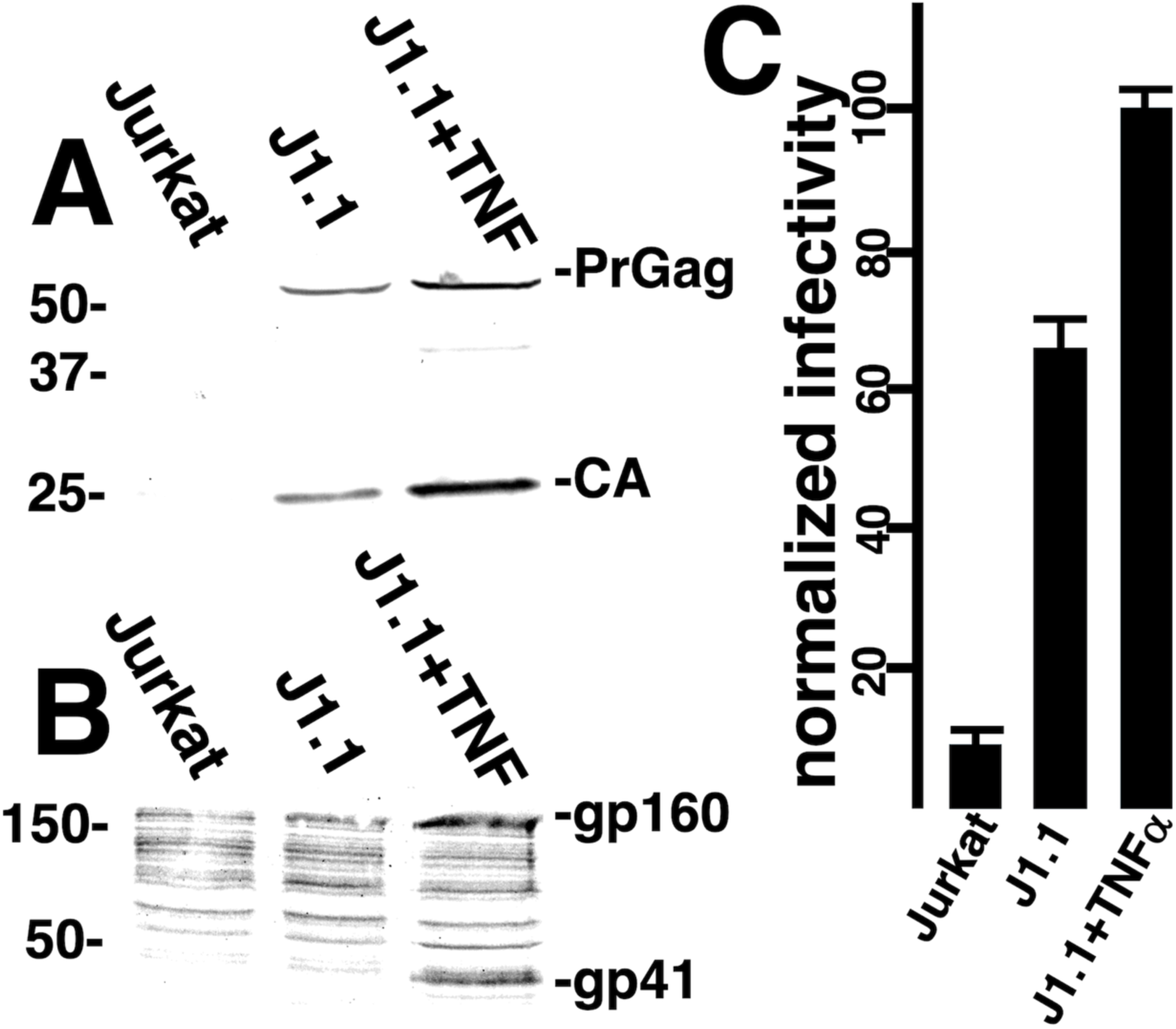
Infection of HIV-1-infected cells. Jurkat T cells, HIV-1-infected J1.1 Jurkat cells, and J1.1 cells induced with tumor necrosis factor alpha (J1.1+TNFα) were subjected to immunoblot detection of Gag proteins **(A)** with an anti-CA antibody, immunoblot detection of Env proteins **(B)** with an anti-gp41(TM) antibody, and infection with a lentivirus vector **(C)** generated by transfection of 293 cells with psPAX2, pLVX-puro-Xho-ATG-βGal, CD4-GPI, and CXCR4-328 plasmids. **(A-B)** Migration positions of PrGag, CA, gp160 and gp41 proteins are indicated, as are molecular weight standards (in kDa) that were electrophoresced in parallel. **(C)** Infectivity results represent normalized transduced βGal activity readings normalized to results for induced J1.1 cells. Averages and standard deviations derive from duplicate (Jurkat, J1.1) or triplicate (J1.1+TNFα) infections, and differences between Jurkat and J1.1, as well as J1.1 versus J1.1+TNFα were highly significant (P<0.001)

### Effects of membrane lipid compositions

As mentioned in the Introduction, HIV-1 particle membranes are highly enriched in PI(4,5)P2, PS, PE, saturated PCs, cholesterol, Cer, SM and HEX (14-24; Figure 8A). Like cellular PMs, the internal leaflets of viral membranes are enriched in PI(4,5)P2, PS and PE lipids, while external leaflets have higher concentrations of SM, HEX, and saturated PC lipids (14-24; Figure 8A). We and others have shown that pertubation of viral cholesterol with AME inhibits replication of Env WT viruses, but not ΔCT viruses (13, 26–28). Additionally, depletion of viral long chain SLs by production of viruses in CerS2-/- cells inhibits both WT and ΔCT virus fusion capabilities, while assembly of a scramblase into virions blocks virus replication (25, 34).

**Figure 8.**
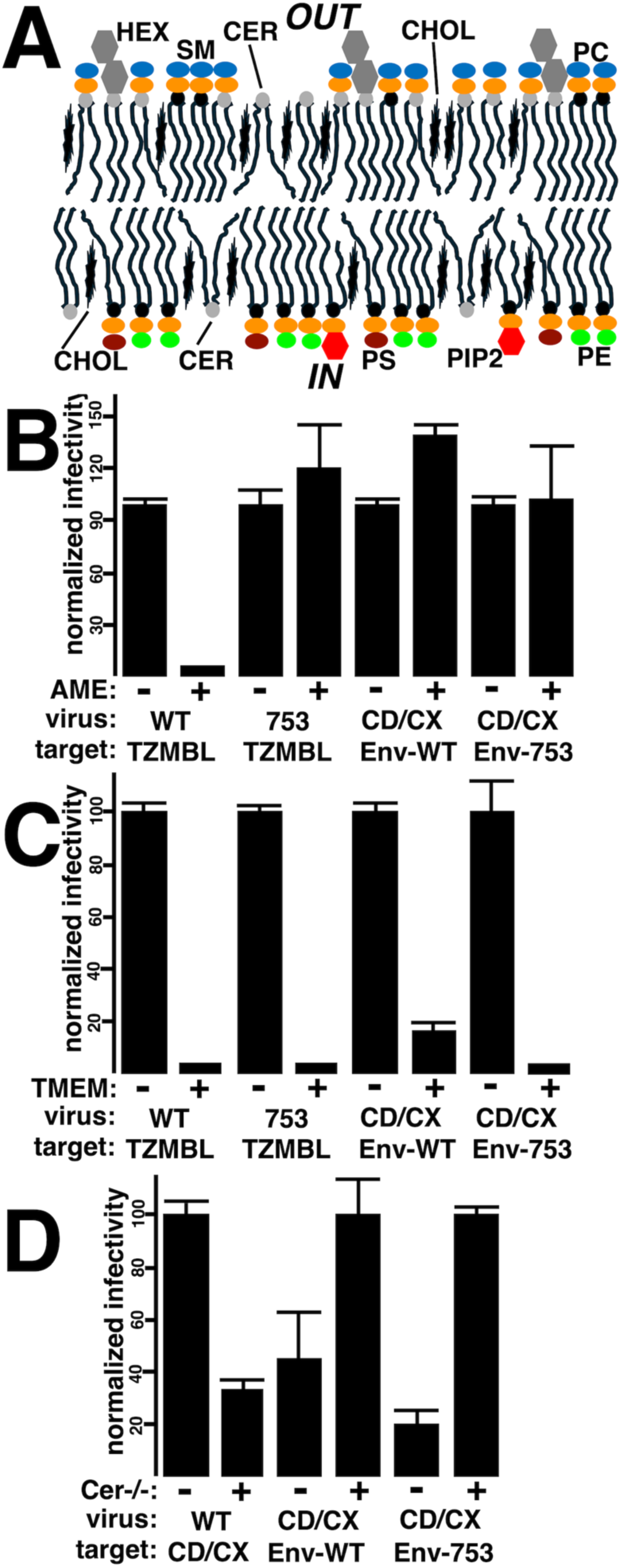
Effects of lipid variations on fusion in reverse infections. **A.** The HIV-1 viral membrane is highly enriched in cholesterol (CHOL), and has substantial amounts of ceramide (CER). The inner leaflets are enriched for phosphatidylserine (PS), phosphatidylinositol-4,5-bisphosphate (PI[4,5]P2; PIP2), and phosphatidylethanolamine (PE); while the outer leaflet is enriched in sphingomyelin (SM), hexosylceramides (HEX), and saturated phosphatidylcholines (PCs). **B-D.** Infectivities were scored via βGal activities, and were normalized either to untreated (-) samples, or for convenience in the panel **D** CD/CX infections, to viruses from CerS-/- cells. In all panels CD/CX viruses refer to lentivirus vectors generated by cotransfection of 293 cells with a GagPol expression vector (psPAX2), a βGal transducing vector (pLVX-puro-Xho-ATG-βGal), as well as CD4-GPI and CXCR4-318 plasmids. Also, in all panels Env-WT and Env-753 targets cells refer to 293 cells transfected to express those proteins. For panels **B** and **C**, WT and 753 viruses are the NL4-3 variants, and were used to infect TZM-bl reporter cells; for panel **D**, WT refers to viruses produced from cells transfected with psPAX2, pLVX-puro-Xho-ATG-βGal, plus a WT Env expression vector, while targets were 293 cells transfected to express the CD4-GPI and CXCR4-318 proteins. In panel **B**, AME refers to treatment of viruses with 10 μM AME prior to infections. In panel **C**, TMEM refers to cotransfection of virus producing cells with (+) or without (-) the mTMEM-GY scramblase expression vector. In panel **D**, CerS2-/- pertains to producing viruses either from 293 cells (-) or from 293-CerS2-/- knockout cells (+). For panels **B-D**, averages and standard deviations are as indicated, and significant differences were as follows: Panel **B** WT/TZMBL, AME -/+, P<0.001; Panel **B** CDCX/Env-WT, AME -/+, P<0.001; Panel **C** all pairs P<0.001; Panel **D** CDCX/Env-WT, CerS2-/-, -/+, P<0.01; all other Panel **D** pairs, P<0.001.

The availability of viruses capable of reverse fusion gave us the unique opportunity to examine how perturbation of receptor plus co-receptor containing membranes might affect HIV-1 replication. For these investigations, we compared how AME, CerS2-/- knockout, and TMEM16F scramblase incorporation affected normal HIV-1 infections versus fusion in reverse infections (Figure 8B-D). Our results with AME pertubation of viral cholesterol are graphed in Figure 8B. As expected (13, 26–28), AME treatment of WT HIV-1 dramatically reduced infection, but did not inhibit infection of Env-753Stop viruses, which behaved similarly to ΔCT viruses. Interestingly, AME treatment of receptor plus co-receptor pseudotyped virions did not impair infectivity, regardless of whether the target cells carried Env-WT or Env-753Stop. These results suggest that the HIV-1 Env fusion machinery can tolerate tampering with cholesterol levels in receptor-plus co-receptor-containing membranes.

Our results with viruses produced from cells expressing the PM-targeted TMEM16F scramblase are shown in Figure 8C. As previously demonstrated (34), WT HIV-1 virions produced in the presence of this TMEM16F variant were essentially blocked for replication. Our observations showing an inhibition of Env-753Stop viruses also are consistent previous results that demonstrated a signficant scramblase inhibition of ΔCT HIV-1 replication (34). Importantly, the TMEM16F scramblase also dramatically reduced fusion in reverse infections for both Env-WT and Env-753Stop target cells. Thus, the HIV-1 Env fusion activity appears to require asymmetric target membranes.

The third lipid variation we tried was a comparison of viruses produced from WT 293 cells as opposed to 293 CerS2-/- knockout cells (Figure 8D). For the WT virus, as seen previously (25), production in CerS2-/- cells reduced viral infectivity 2- to 3-fold. In contrast, and surprisingly, production in CerS2-/- cells increased the efficiency of fusion in reverse, implying that the presence of long chain SLs in receptor plus co-receptor membranes may have a negative effect on HIV-1 Env activities. These results are discussed in more detail below.

## DISCUSSION

Ordinarily, a small number of trimeric HIV-1 Env proteins are incorporated into HIV-1 particles, and must perform their receptor-binding, co-receptor-binding, and fusion duties within the atypical viral lipid membrane environment that features high levels of PI(4,5)P2, PS, PE, saturated PCs, cholesterol and sphingolipids (1-9, 14-24; Figure 8A). However, it’s long been known that Env trimers on the surfaces of infected cells can mediate cell-cell fusion with neighbor receptor-plus co-receptor-expressing cells, resulting in syncytia formation (35–39). Using a split GFP reporter system (Figure 1A), we examined the capacities of different CD4 receptor plus CXCR4 co-receptor combinations to participate in cell-cell fusion reactions with cells expressing either WT or C-terminally truncated (753Stop) Env proteins. Among our observations were that CD4-GPI and truncated CXCR4-318 proteins were competent for fusion reactions, and that fusion efficiency with Env-753Stop was significantly higher than with Env-WT. Because Env CTs can influence Env fusogenic properties (13, 27–28, 70–74), it is possible that an increase in Env-753Stop fusogenicity could account for some of the efficiency difference observed between WT- and 753Stop-directed cell-cell fusion. However, the ten-fold difference in cell-cell fusion efficiencies directly matched the cell-surface expression differences of the two Env proteins (Figures 2-3), which suggests that Env surface expression is responsible for the disparity.

Based on previous observations of fusion in reverse (40–48), we tested CD4 and CXCR4 variants for their abilities to be assembled into HIV-1-based lentivirus vectors and accomplish fusion in reverse infections. Our results clearly demonstrate that this depends on both a receptor and co-receptor (Figure 5), and on the fusion function of Env (Figure 4). In addition to the truncated CXCR4-318 variant, other CXCR4 variants also were capable of mediating infection, including WT CXCR4 (Figure 5). Considering previous results (40–47, 57–58), this is not surprising. In contrast, while WT CD4 facilitated cell-cell fusion reactions (Figure 2), it was deficient for fusion in reverse reactions (Figure 5). Given that CD4 and CD4-GPI virion incorporation levels appeared similar (Figure 6), our results imply that in the context of HIV-1 membranes, CD4-GPI proteins are more active than WT CD4 proteins either in Env binding or collaboration with CXCR4 proteins in directing fusion. In this regard, it is pertinent to mention that a previous study (45) did not see a difference in CD4- and CD4-GPI-directed fusion in reverse. We speculate that this could be due to specific differences in vectors, protocols, and detection methods employed, and note that viral CD4 and CD4-GPI levels were not monitored in those investigations (45).

When we extended our analyses to include HIV-1-infected cells, we observed that latently infected Jurkat J1.1 cells (65–69) were specifically infected, and that induction of HIV-1 expression increased the efficiency of infection (Figure 7). These results again (40–47) demonstrate that a fusion in reverse approach permits the specific transduction of cells harboring HIV-1, albeit with the caveat that the efficiency depends greatly on Env surface expression levels (Figures 3, 4, 7; 42). In addition to CXCR4-tropic envelopes, others have shown that CCR5-tropic Envs (75–77) can be targeted (42–43, 47). In this regard, it also is tempting to speculate that some Env-binding nanobodies (78) may precisely replicate co-receptor properties and facilitate fusion in reverse, but no evidence in favor of this possibility is currently available.

With respect to other applications of the HIV-1 fusion in reverse approach, we’ve demonstrated that it can allow the examination of factors that determine whether membranes carrying receptors and co-receptors are capable of Env-directed fusion. Interestingly, we did not find that AME perturbation of membrane cholesterol affected infection (13, 26–28), but that assembly of virions in the presence of the PM-targeted TMEM16F scramblase (34) effectively killed the process (Figure 8). This observation strongly suggests that target membrane leaflet asymmetry is essential for efficient Env-directed fusion. We additionally observed that rather than inhibiting infectivity (25), expression of fusion in reverse virions in cells with a deficiency in very long chain SLs increased infectivity (Figure 8). Despite these intriguing results, it is pertinent to note that experiments performed with GPI-anchored CD4 may or may not reflect what might occur with WT CD4. Nevertheless, we believe that the HIV-1 fusion in reverse system will prove useful in the dissection of other factors that regulate HIV-1 Env-driven membrane fusion.

## MATERIALS AND METHODS

### Recombinant DNA constructs

The lentivirus GagPol packaging plasmid (psPAX2), the GFP expression plasmids (GFP-PS, GFP-NoPS, GFP-C1-PLCdelta-PH, pEGFP-GFP11-ClathrinLightChain, pcDNA3.1-GFP[1-10]), the VSV G expression vector pMD.G, the CD4 expression plasmid pCAGGS-CD4-Myc, and the CXCR4 expression plasmid pcDNA3-CXCR4 all were obtained from Addgene, and were the gifts of Drs. Didier Trono, Jennifer Doudna, Tobias Meyer, Bo Huang, Simon Davis, Erik Procko and Jacob Yount (53, 62, 79–83). The TMEM16F scramblase expression vector, mTMEM-GY, was kindly provided by Dr. Walther Mothes (34). The mammalian expression plasmid pcDNA3.1 (Invitrogen), the HIV-1 envelope expression vector pSVIII-Env (from the NIH HIV Reagent Program), the pLVX-puro-Xho-ATG-βGal β-galactosidase lentivirus vector, and the wild type (WT) and 753Stop HIV-1 NL4-3 proviral constructs have been described previously (13, 48, 84–85). The pMDG-NL43-Env-WT expression vector was created from the pMD.G backbone (81). The pMD.G EcoRI site at nt 3774 first was killed by insertion of an A residue after nt 3775, giving the sequence 5’ tctaga**a**attc 3’, where the 5’ nt corresponds to pMD.G nt 3771, the 3’ nt corresponds to nt 3880, and the inserted nt is in bold. After that, the VSV G encoding EcoRI fragment was replaced with an EcoRI fragment containing the NL4-3 envelope protein coding region from the NL4-3 nt 6153 SspI site to the nt 8887 XhoI site. The 5’ juncture sequence is 5’ *gaattc*aagc ttgtcgacct gcaggcatgc aagcttcag**a** tt 3’, where the EcoRI site is italicized and NL4-3 nt 6156 is in bold. The 3’ juncture sequence is 5’ **ctcgag** *gaattc* 3’, where the NL4-3 XhoI site is in bold and the EcoRI site is italicized. The pMDG-NL43-Env-753Stop construct is identical to pMDG-NL43-Env-WT except that it carries a termination codon after HIV-1 Env residue 753, yielding the reading frame between NL4-3 residues 8465 and 8476 5’ gga tcc TAA gca 3’ (85). The pCAGGS-CD4-GPI construct was generated from pCAGGS-CD4-Myc by replacement of the CD4 transmembrane and cytoplasmic domains with the GPI anchor from the human placental alkaline phosphatase (PLAP) protein (48, 86–87). To do so, CD4 was truncated at codon 393 and fused to the AscI-XhoI PLAP GPI fragment described in the construction of the previously described pMDG-Nano-GPI plasmid (48). The reading frame at the CD4-GPI juncture is as follows, where the bold codon indicates CD4 codon 393: 5’ tgg tcc acc **ccg** gcg cgc cag 3’. For generation of the pcDNA3-CXCR4-309Stop and pcDNA3-CXCR4-318Stop cytoplasmic truncations of CXCR4, the parental pcDNA-CXCR4 plasmid was modified. The reading frame at the CXCR4 C-terminus of pcDNA3-CXCR4-309Stop is 5’ gcc aaa **ttc** TGA ggc ctc gag 3’, where the codon for residue 309 is in bold, and the stop codon is capitalized. The reading frame at the CXCR4 C-terminus of pcDNA3-CXCR4-318Stop is 5’ ctc **acc** TGA ccc ggg ctc gag 3’, where the codon for residue 318 is in bold, and the stop codon is capitalized. The CXCR4 variants with C-terminal fusions to the phospholipase C delta pleckstrin homology domain (PLCδPH) were based on a variant of GFP-C1-PLCdelta-PH (62–64) in which the GFP N-terminal coding region from the NheI site to the BamHI site at the N-terminus of the PLCδPH domain was replaced with CXCR4 fragments. For CXCR4-309-PH the CXCR4-PLCδPH juncture sequence is 5’ gcc aaa **ttc** ggg gga ggc ggc gga tcc ATG 3’, where the codon in bold encodes CXCR4 residue 309, and the capitalized codon is the start of the PLCδPH. For CXCR4-318-PH the CXCR4-PLCδPH juncture sequence is 5’ ctc **acc** tgg gaa ttc ggg gga ggc ggc gga tcc ATG 3’, where the codon in bold encodes CXCR4 residue 318, and the capitalized codon is the start of the PLCδPH. Our split GFP partners also represent fusions to the PLCδPH. For pGFP-H11-PLCdeltaPH, where the eleventh helix of GFP is fused to a C-terminal PLCδPH, the pEGFP-GFP11-ClathrinLightChain (53) plasmid EcoRI-HpaI clathrin-encoding fragment was replaced with the PLCδPH EcoRI-HpaI fragment from GFP-C1-PLCdelta-PH. The resulting GFP helix 11-PLCδPH juncture reading frame is 5’ **aca** ggc gga tcc gga ctc aga tct cga gct caa gct tcg aat tct gca gtc gac ggt acc gcg ggc ccg gga tcc ATG 3’, where the codon in bold represents the last residue of GFP helix 11 and the capitalized codon is the start of the PLCδPH. For pcDNA3.1-PLCdeltaPH-GFP-H1-10, the PLCδPH coding region was inserted in frame and upstream of the GFP helix 1-10 sequence in pcDNA3.1-GFP(1–10) (53). The reading frame at the PLCδPH N-terminus is 5’ *gct agc* gcc acc **atg** gcg gcc gcg gga tcc ATG 3’, where the italicized sequence is the pcDNA3.1 NheI site, the capitalized sequence is the PLCδPH start codon, and the bold sequence is an alternative in frame start codon. The PLCδPH-GFP juncture sequence is 5’ aag gag **ctc** ggg ata tct agc aag ctt ccc ggg ctc gag gaa ttc gtt aac aga tct ATG 3’, where the bold codon indicates a PLCδPH C-terminal residue corresponding to residue 433 of the EGFP-C1-PLCdeltaPH fusion protein and the ATG indicates the beginning of the GFP helix 1-10 sequence.

### Cell culture

Human embryonic kidney 293T cells (293; 88) were obtained from the American Type Culture Collection (ATCC), HIV-1 reporter TZM-bl cells (89–91) were from the National Institutes of Health (NIH) AIDS Reagent Program, and 293-CerS2-/- (CerS2-/-) knockout cells, which carry premature stop codons after 63 residues of both CerS2 alleles, were obtained from Dr. Antony H. Futerman (25, 92). The Jurkat T cell line (93) was from the BEI Resources HIV Reagent Program, as was the J1.1 cell line, which is an HIV-1-infected Jurkat derivative (65–69). Cells were routinely were grown in humidified 5% carbon dioxide air at 37°C in Dulbecco’s Modified Eagle’s Media (DMEM; 293 and CerS2-/-) or Roswell Park Memorial Institute (RPMI) media, both of which were supplemented with 10% fetal bovine sera (FBS) plus 10 mM Hepes, pH 7.3, penicillin and streptomycin (13, 25, 48).

Transfections of cells were performed using polyethyleneimine (PEI) as described previously (13, 25, 48, 94), and typically employed 24 μg total DNA for 5 million cells on 10 cm plates, or 6 μg total DNA for 1 million cells on polylysine-coated (0.01%; Sigma P4707) coverslips (Fisher 22×22-1.0) in six well plates with 35 mm well diameters. Unless otherwise noted, in cases where cells were cotransfected with multiple plasmids, equal DNA amounts in weight for each plasmid were used, to a total of 24 μg. When infectious NL4-3 viruses were scored for infectivity, TZM-bl cells were seeded onto wells of six well plates (1 million cells per well) 24 h before infections. For the generation of 293-derived target cells to be infected, cells were seeded one day post-transfection into six well plate wells at one million cells per well, and grown an additional two days prior to infection. For cells to be processed for immunofluorescence, cells transfected on coverslips were refed one day post-transfection and processed two days later. Alternatively, cells on 10 cm plates were split onto polylysine-coated coverslips one day post-transfection (1 million cells per coverslip) and processed two days later. A third approach was employed for cell-cell fusion assays. In this case, 24 h after separately transfecting cells on 10 cm plates, cells were co-seeded onto polylysine-coated coverslips at 1 million cells per cell fusion partner, and incubated an additional 48 prior to processing for microscopy.

For the production of NL4-3 virus stocks and pseudotyped lentivirus vectors, transfected 293 or CerS2-/- cells were refed one day post-transfection, and cells and virus-containing media supernatants were collected at 3 days post transfection. Cell samples for protein analyses were pelleted and frozen at -80°C prior to use. Viral samples were filtered through 0.45 micron filters, and either used immediately, or stored at -80°C. In some cases, viral samples were processed for protein analyses. For this, filtered virus-containing media samples were concentrated by centrifugation (1 h at 197,000 x g; 35,000 rpm, Beckman SW41 rotor) through 2 ml cushions of 20% or 30% sucrose in phosphate-buffered saline (PBS; 9.5 mM sodium potassium phosphate [pH 7.4], 137 mM NaCl, 2.7 mM KCl), suspended in 0.1 ml PBS, and stored at -80° prior to use.

Viral infections were performed with cells seeded in six well plates and filtered viruses. Typically, cells were infected 3-4 h with 1 ml viral stocks at 37°C, supplemented with 1 ml media, and incubated an additional 72 h prior to collection for β-galactosidase (βGal) assays, which were performed as described previously (48, 95). Briefly, media on cells in six well plate wells (35 mm diameter) were removed, and cells were scraped into 1.0 ml PBS and pelleted. Cell pellets were suspended in 150 μl PBS containing 0.1% sodium dodecyl sulfate (SDS), vortexed, supplemented with 600 μl PM-2 buffer (33 mM NaH2PO4, 66 mM Na2HPO4, 2 mM MgSO4, 0.1 mM MnCl2, 40 mM β-mercaptoethanol [BME]), vortexed, supplemented with 150 μl 4 mg/ml 2-nitrophenyl-β-d-galactopyranoside (ONPG) in PM-2 buffer and incubated at 37°C. Reactions were stopped by addition of 375 μl 1 M Na2CO3 and flash freezing on dry ice powder. Samples then were thawed, and 420 nm light absorbances were read spectrophotometrically to calculate β-galactosidase activities (1 unit = 1 nMole ONPG hydrolyzed per minute = 420 nm absorbance x 285/minutes of incubation) as a measure of infectivity (48, 95). To calculate significance values, means and standard deviations from multiple experiments were determined and converted to Z values, which were used to calculate probabilities.

In addition to the above infection protocol, three variations also were employed. When viruses were treated with amphotericin B methyl ester (AME; kindly provided by the National Cancer Institute Developmental Therapeutics Program [NCI DTP; NCI #252624] and used after minimum freeze-thaw cycles to avoid degradation), they were incubated for 2 h at 37°C in the presence of 10 μM AME before infections (13, 26–28). For testing of the effects of T-20 (Enfuvirtide; BEI Resources HIV Reagent Program; 59-61) on infections, cells were infected 24 h in 1 ml media plus one ml virus plus either 10 μM T-20 in dimethylsulfoxide (DMSO) or DMSO (final 1%). After 24 h, infection mixes were removed, cells were washed once with media, refed with media without T-20 or DMSO, and incubated an addtional 48 h, prior to assay. Infections of non-adherent Jurkat, and J1.1 cells also employed a slightly modified approach. Here, cells either were mock treated or treated for 48 h with 10 ng/ml tumor necrosis factor alpha (TNF-α; Stemcell Technologies #78068.1). After this treatment, cells were infected for 72 h at 37°C in incubations containing 1 ml media, 1 ml virus and 3 million cells in the presence or absence of 10 ng/ml TNF-α. Following infections, pelleted cells were processed for βGal assays as described above.

### Microscopy

For direct viewing of cell-cell fusion incubations, cells on coverslips were washed with PBS, fixed 20 min at room temperature in 4% paraformaldehyde (PFA; Sigma) in PBS, washed three times in PBS, incubated 5 min at room temperature with 200 ng/ml 4’,6-diamidino-2-phenylindole (DAPI; Sigma) in PBS, washed three times in PBS, and mounted onto microscope slides in Fluoro-G (Thermo-Fisher) mounting medium. For indirect immunofluorescent detection of cell surface Env proteins, cells on coverslips were washed and fixed as described above and then processed for immunofluorescence detection as described previously (48, 64, 96), except excluding the Triton X-100 permeabilization step. Primary antibodies were human anti-gp120 VRC03 and 2G12, respectively used at 1.67 μg/ml and 0.67 μg/ml; and the secondary antibody was an AlexaFluor594-conjugated goat anti-human IgG(H+L) antibody (Invitrogen A11014), used at 1:1000 (2 μg/ml).

Fluorescence viewing was on a Keyence BZ-X710 fluorescence microscope using either Plan Fluor 20x or Plan Fluor 40x (oil) lenses and the following filter sets: DAPI, excitation 360 nm /40 nm slitwidth, emission 460 nm/50 nm slitwidth; GFP, excitation 470 nm/40 nm slitwidth, emission 525 nm/50 nm slitwidth; AlexaFluor594, excitation 560 nm/ 40 nm slitwidth, emission, 630 nm/75 nm slitwidth. All imaging employed 100% gain settings, and samples to be compared used identical exposure settings, unless otherwise noted.

For cell-cell syncytia counting of each sample, numbers of GFP+ syncytia per five 0.4 mm^2^ fields were counted and averaged. For determination of Env+ cell percentages, red-staining cell counts and total cell counts from a minimum of nine images for each sample were used in calculations. For comparison of cell brightnesses, background-subtracted gray scale brightness values were determined for boxed Env+ cells using (Image J [Fiji]; 97) software. Raw brightness values then were divided by image exposure times to yield final brightness values. Note that for Env-753Stop-expressing cells, exposure times typically were 10 ms, while for Env-WT-expressing cells, exposure times were 67-100 ms. Note also that brightness value averages and standard deviations were calculated from a minimum of 14 cells for the VRC03 primary antibody, and a minimum of 36 cells for the 2G12 primary antibody.

### Protein analysis

For analysis of concentrated virus samples, pelleted and resuspended viral samples were mixed with an equal volume of 2x sample buffer (12.5 mM Tris-HCl [pH 6.8], 2% SDS, 20% glycerol, 0.25% bromphenol blue) plus 5% β-mercaptoethanol (BME), and frozen at - 80°C. Cell samples for protein analysis were prepared by collecting cells in PBS, pelleting 20% of the cell sample, suspension in 50 μl IPB (20 mM Tris-HCl [pH 7.5], 150 mM NaCl, 1 mM ethylenediamine tetraacetic acid [EDTA], 0.1% sodium dodecyl sulfate [SDS], 0.5% sodium deoxycholate, 1.0% TritonX-100, 0.02% sodium acetate), vortexing, incubation on ice for 5 min, pelleting 15 min at 13,000 x g to remove insoluble debris, mixing with 50 μl 2x sample buffer plus 0.1 volume of BME, and stored frozen at -80°C prior to analysis.

Samples for protein analysis were subjected to SDS-polyacrylamide gel electrophoresis (SDS-PAGE) as described previously (13, 25, 48, 64, 95–96), except that samples were heated only to 37°C before electrophoresis, and acrylamide concentrations were either 10% or 7.5%. Typically 10% of total viral samples or 4% of total cell samples were subjected to electrophoresis in parallel with molecular weight size standards (Bio-Rad). After SDS-PAGE fractionation, proteins were electroblotted and immunoblotted following previously described methods (13, 25, 48, 64, 95–96). Primary antibodies employed were as follows: mouse anti-HIV-CA hybridoma media (Hy183, kindly provided by Dr. Bruce Chesebro) at 1:15; mouse anti-gp41(CT) hybridoma media (Chessie 8, recognizing CT residues 727-732 [PDRPEG], from the NIH AIDS Reagent Program) at 1:15; rabbit anti-CD4 D2E6M monoclonal antibody (Cell Signaling Technology, 93518T), used at 1:2000; polyclonal rabbit anti-CXCR4 antibody (ProSci, #1009), used at 1:2000. Secondary reagents were alkaline phosphatase-conjugated anti-mouse, anti-human, or anti-rabbit IgG antibodies (Promega) used at 1:15,000. Color reactions for visualization of antibody-bound bands employed nitrobluetetrazolium plus 5-bromo-4-chloro-3-indolyl phosphate in AP buffer (1000 mM Tris-hydrochloride [pH 9.5], 100 mM NaCl, 5 mM MgCl2; 13, 25, 48, 64, 95-96). For quantitation, immunoblots were air-dried an scanned using an Epson Perfection V600 scanner, and band intensities of scanned TIFF images were determined using NIH Image J software (97).

## ACKNOWLEDGMENTS

We are grateful to Dr. Antony Futerman for providing the CerS2-/- cells, and to Addgene, the NIH AIDS Reagent Program, BEI Resources and Didier Trono, Jennifer Doudna, Tobias Meyer, Bo Huang, Simon Davis, Erik Procko, and Jacob Yount for making reagents and recombinant DNA constructs available. EB and FT also are thankful for funding from the National Institutes of Health through grants RO1 AI152579 (EB) and RO1 AI141549 (FT).

## AUTHOR CONTRIBUTIONS

EB and AA conceptualized, performed, and analyzed the experiments and wrote the manuscript. EB also was involved in funding acquisition and providing resources. RLB was involved in methodology development and performing experiments. FT provided funding and resources, analyzed data, and helped in manuscript revision.

